# On Miller’s rule for the brain working memory, or why human memory is so short

**DOI:** 10.1101/035014

**Authors:** E. Z. Meilikhov, R. M. Farzetdinova

## Abstract

Working memory is a cognitive construct that describes how information can be maintained in brain for a limited period of time, while concurrent processing is also performed. We present a simple model that accounts for working memory span and explains the origin of the cognitive Miller’s rule (Magical Number Seven).

---

Working memory is one of basic concepts of cognitive psychology. In the framework of various models^1^ it is understood to be the complex of structures and processes that provide operational (relatively short-term) information integrity in the course of time sufficient to use it in cognitive processing.

In the simplest model, memory consists of only three components: sensory registers plus short-term and long-term information repositories^2^. In that model, the short-time repository is a passive structure for operational storage of the verbal and iconic material (with phonological loop and visual-spatial scratch-pad) only. Herewith, the repetition process prevents decaying the information stored.

In more elaborated model (for instance, that of Baddeley^3^), processes of information handling are equally important along with the storage. In that model, there is the central control instance (the so called central executive) whose functions are coordinating information processing and controlling the storage system.

Numerous experiments and life experience provide support for the finite capacity of the short-term information depot^4,5^. Various authors estimate the latter by elements’ number from 3 to 7 ± 2 (“magic Miller’s number”)^6,7^. Such a high scatter is to a great extent connected with differences of experimental procedures taken as a basis of determining the absolute value of the working memory capacity. For example, in the traditional method, which dates back to the end of nineteen century^8^ and consists in presenting arrays of digits, they estimate the so called digit span of memory. Different versions of that technique are up to now used (see, for instance, the Wechsler test^9^) applying towards estimating intellectual abilities.

More complex tests include stages of cognitive information processing, which obstruct (or suppress) repetition mechanisms meant for extending information storage time in working memory. It is clear that such more complicated techniques should lead to the underestimated span of the working memory (compared to the classic Miller’s result). Hence, the possibility to “whip up” decaying memory (similarly to whipping a tired horse) is of critical importance for extending time of information storage in working memory.

Attempts are known to deduce the Miler’s rule on the basis of some speculative ideas concerning the manner of writing and storing information in the brain neuronal net (cf., for instance,^10^). Though the accurate form of the Miller’s rule is significant (for example, for WEB designers), no less important to understand (proceeding from some experimental facts) why the Miller’s number ranges within the relatively narrow interval 3–9 (i.e., on the order of 10) and, definitely, does not equal to 10^3^ or 10^2^.

The phonological loop associated with the process of articulatory repetitions is the critically important construct that defines the confinement and reproduction of information^11^. Without the loop, information (a “trace” in the working memory) decays over a period of about a few tens of seconds. Those traces could be refreshed by the vocal or mental articulatory replay which, in fact, is the phonological loop. With high enough frequency of using identical loops, concerning some specific information, the latter could be preserved “eternally”. As for the number of such eternally stored information elements (that is, the Miller’s number), it is defined, obviously, by the relation between typical times of decaying and renovating the information (see below).

Let us pass now to the mathematical modeling of that process. To describe the grade of the information integrity quantitatively, we introduce the “order parameter” *x*, that could take values in the range from 0 up to 1.

The value *x* =1 corresponds to the maximum unharmed and readily accessible information in the working memory, while the value *x* = 0 associates with the disappeared (or, anyhow, inaccessible in a “reasonable” time) information. Intermediate values of the parameter *x* relate to the information which is, to any extent, accessible for extracting. The higher is the parameter *x*, the faster and more accurate that information is extracted from memory. From the practical point of view, all low enough values of the order parameter (*x < x_c_*) are the same: extracting zero information in infinite time (at *x* = 0) is equivalent to obtaining a small part of information in finite but very long time (on order of one hour, for example). By this is meant that there is such a critical value *x_c_* of the order parameter, which divides the whole of information stored in memory into two sorts – accessible (*x > x_c_*) and practically inaccessible (*x < x_c_*) ones. The critical value *x_c_* depends on the specific organization (“design”) of the storage brain structure and could not be unambiguously defined today. In that situation, there are two possibilities: 1) to hope that in the framework of the considered mathematical model (see below) the value *x_c_* influences final conclusions slightly enough, or 2) to use some general considerations concerning the possible *x_c_*-value, basing on some general ideas about storing and retrieving information.

Presenting stimulus results in a response – modifying the state of large number of neurons^12,13^. The assembly of such “excited” neurons makes the net object (array of sites connected with bonds), or a graph. It is reasonable to suggest that information, coded by that graph, is survived, in more or less full measure, until the graph connectivity is preserved, i.e., there are paths along chains of relevant neurons between every sites of the neuron graph. Existence or non-existence of such a connectivity is the typical problem of the percolation theory^14^.

It is known that the connectivity condition for the graph with a number of sites depends on it’s geometrical properties and, particularly, upon if the graph is regular (makes the regular lattice), random or presents somewhat intermediate – for example, the so called Small World network^15^. In the case considered, such graphs are, likely, not regular. Thus, we could use known results from the percolation theory according to which the connectivity of random graphs is violated when the fraction of non-broken bonds (or non-removed sites) amounts to ~ 20 – 50%^14^. That gives reason to assume, for instance, *x_c_* ≈ 0.35 (that corresponds to the middle of the range indicated).

Thus, we characterize the memorizing level of some element by the parameter *x*, taking values between 0 and 1 with values *x > x_c_* corresponding to the accessible information, and *x < x_c_* – to the inaccessible (disappeared) information. Without using the phonological loop, the information “written” in working memory decays with time. According to the empirical Jost’s low (older information is forgotten more slowly), that decay is exponential^16,17^:

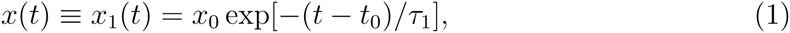

where *t* is current time, *x*_0_ is the order parameter at the moment *t* = *t*_0_. The basic parameter in that relation is the characteristic time *τ*_1_ of information storing in shortterm memory, whose typical values amount to ~ 30 s^17^, ~ 20 s^18,19^, 15 – 30 s^20,21^. If one assume *x*_0_ = 1 (that corresponds to the maximum memory trace) and *t*_0_ = 0 (i.e., if one measures time from the moment when *x* = *x*_0_ = 1), then according to (1) in the time *τ*_1_ the order parameter drops down from *x* = 1 to *x* ≈ 0.37 ≈ *x_c_*. In other words, the time *τ*_1_ is that of dark oblivion, and if we do not like to allow that we have to use some remedy which prevents forgetting.

Such a remedy is, as well known, the phonological loop which “refreshes” effectively memory traces.

Formal consequence of using phonological loop is the growth of the order parameter *x*. The longer the loop “works”, the higher that parameter grows. However, that growth is limited above by the maximum value *x* = 1. Therefore, by analogy with the 2nd Jost low (the low of verbal learning)^16,17^, the process of refreshing information could be described by the relation

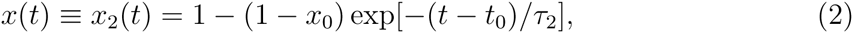

where, again, *t* is the current time, *x*_0_ is the order parameter at the moment *t* = *t*_0_, *τ*_2_ is a characteristic time of restoring memory. If one assumes *x*_0_ = 0.4 (that is close to the threshold value *x_c_* = 0.35 taken above), then in the time *t − t_0_* = *τ*_2_ the order parameter grows up to the value *x* ≈ 0.78 which is significantly bigger than the critical value, and in the time *t − t*_0_ = 2*τ*_2_ it reaches the value *x ≈* 0.92 that corresponds to practically complete recovery of the pattern in memory. As for the duration of the “restoration” time *τ*_2_, it is comparable with the articulatory time and amounts *τ*_2_ ~ 1 − 3 s.

Thus, to hold possibly bigger number of elements in memory one should *consequently* (because parallel using more then one phonological loops is impossible) refresh the information relating to each of those elements, providing the values of parameters *x_i_* (*i* = 1, 2,…) to be always higher than the threshold value *x_c_*. That resembles juggling – every thing should be insured against falling by on-time catching and skiing. World records of juggling (registered in The Guinness Book of Records) are, for example, 10 and 8 for balls and clubs, correspondingly (juggling within several seconds) and 5 (that in about 10 minutes). Curious, the latter figure is fairly get through the extended Miller’s rule. Perhaps, that coincidence is not accidental.

With using phonological loop, one passes consequently from articulating a given element to that of the next one. Assume, that at each of those stages the relevant order parameter *x_i_* is restored from a value near to *x_c_* up to the value *x ≈* 0.9. This, as we have seen, takes time near to 2*τ*_2_. On the other hand, refreshed information degenerates down to the critical level in the time *τ*_1_. It is this time interval that is offered for “serving” other decaying images. It is obvious, that the number of images in memory, which one could support with the described technique, equals to the ratio

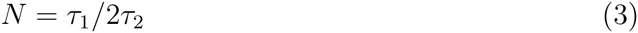

of the decaying time to that of restoring an image in working memory. Making use of cited above values *τ*_1_ ~ 30 s and *τ*_2_ ~ 2 s, one finds *N ≈* 7, that agrees with the classical result of Miller.

How critical is assuming the threshold value *x_c_*? Repeating former calculations for other values of *x_c_* within the interval 0.2 < *x_c_ <* 0.5, one finds *N ≈* 9 for *x_c_* = 0.2 and *N ≈* 5 for *x_c_* = 0.5, that agree with boundaries of the confidence interval of Miller (*N* = 7 ± 2). Hence, the choice of the threshold value *x_c_* is not crucial and, what is more, it helps to understand possible nature of scattering results in Miller’s experiments.

In fact, the result (3) is the consequence of the qualitative “time-based resource-sharing” conception ^22,23^ assuming the interplay between temporal decay and refreshment of information in working memory. In our model, the working memory span is connected not with the size of the physical space of some brain fields or the density of neuronal nets, but, more, with relaxation characteristics – the slower information decays in working memory and the sooner it is re-established with phonological loops, the higher the memory volume.

Suggested model also explains naturally the so called word-length effect: the number of elements bearing in memory depends on time required to articulate corresponding words^24,25^. In fact, from Eq. (3) it follows *N* ∝ 1/*τ*_2_, but for long words *τ*_2_ is longer, that just explains the mentioned effect.

In the framework of the considered model, one could also explain why chunking, i.e. grouping separate elements in units of higher association level, does not change the number of remembered images^6^. In this regard, it is sufficient to assume that times *τ*_1_*, τ*_2_ are proportional to the information volume of a “chunk”, so that the complex images are restored and relaxed longer than simple ones. Then the ratio *τ*_1_/*τ*_2_ of relaxation times keeps to be unchanged and, hence, the number of remembered elements does not vary. That just explains the phenomenon of the Miller’s wallet (there is seven coins in wallet, independent of their face values).

Surely, we do not have to attach great importance to the numerical proximity of obtained estimates and experimental values because it depends, in great extent, on choosing specific values of parameters *τ*_1_, *τ*_2_, and in view of the apparent roughness of the suggested model, as well. However, we hope that the model gains an insight into the most important features of the phenomenon, termed the working memory.

Authors are grateful to Prof. B.M. Velichkovsky for valuable discussions.

